# Trait-Dependent and Site-Specific Effects of Foliar IAA and Kinetin on Two Chickpea Varieties Grown Under Contrasting Conditions

**DOI:** 10.64898/2026.08.29.747965

**Authors:** Zoubir Kebaili, Abdelouahab Yahia, Ayache Boultif, Adlene Benyamina, Warda Azabi, Sirid Remache, Amar Zellagui, Merzoug Benahmed, Noureddine Gherraf

## Abstract

Chickpea (Cicer arietinum L.) yield in Algeria still falls short of domestic demand, and low-cost agronomic tools such as plant growth regulators are one of the few levers producers can adjust without heavy investment. We tested the separate and combined effects of foliar-applied indole-3-acetic acid (IAA) and kinetin (0, 10, and 20 mg/L) on two chickpea varieties, FLIP 84-92 and ILC 32-79, grown side by side in the field at the ITGC station of El Khroub and in pots at the Institute of Natural Sciences of Oum El Bouaghi during the 2024/2025 season. Nine hormone treatments were applied to each variety at each site, with three replicates apiece. No single treatment came out on top across the board. A20+K20 gave the largest pot leaf area at both sampling dates (5.67 and 7.20 cm2) and the highest field dry matter weight (95.21 g), yet A10+K10 produced the largest field leaf area after the first spray (16.47 cm2) and the highest pot dry matter weight (6.12 g). Kinetin alone at 20 mg/L (K20) gave the most field pods (21.28) and the most field leaves after the first spray (115.41), while A10+K20 gave the heaviest pot grains (100-grain weight of 21.16 g). Field and pot means are not directly comparable given how different the two growing environments and sampling routines were. Between varieties, FLIP 84-92 germinated better (86% vs. 83%) and produced heavier grains, while ILC 32-79 grew taller stems. Because the manuscript we worked from supplied only treatment means, with no replicate-level data or variance estimates, we report these as descriptive numerical differences rather than statistically tested effects. Read that way, the pattern that emerges is that IAA and kinetin responses are trait- and environment-specific rather than uniformly additive, and a properly replicated factorial analysis will be needed before any interaction or synergy between the two hormones can be claimed.

## 1. Introduction

Grain legumes remain one of the most efficient sources of plant protein available to farmers working nitrogen-poor soils, since their capacity for symbiotic nitrogen fixation cuts fertilizer costs substantially. Chickpea (Cicer arietinum L.) is particularly well suited to this niche: a deep, well-developed root system gives it real drought tolerance, which is why it keeps showing up in dryland rotation systems across semi-arid regions (Cubero, 1987; Duke, 1981; Van der Maesen, 1987). Flowering and early reproductive development make up the critical window for chickpea yield, so whatever management choices are made during this stage tend to carry disproportionate weight for the final harvest (Lake and Sadras, 2014).

Algerian chickpea production has long lagged behind domestic consumption, a shortfall usually attributed to the climatic and agronomic constraints of growing the crop under semi-arid conditions. Specific production and deficit figures reported in earlier drafts of this work could not be traced to a verifiable source and year, so we have left them out here rather than repeat an unconfirmed number.

Plant growth regulators are one of the few agronomic levers available to address this gap, and although they have been studied in chickpea for decades, they remain underused in practice; earlier trials applying growth regulators to chickpea have reported measurable gains in nodulation and yield, which is part of the rationale for testing IAA and kinetin again here (Fatima et al., 2008). IAA governs cell elongation and apical dominance, while kinetin drives cell division, slows leaf senescence, and redirects nutrients toward actively growing tissue (Heller, 1985; Guignard, 2000; Mazliak, 1982). The two hormones do not simply add up when applied together: auxin and cytokinin signal through pathways that overlap heavily even though they originate separately, acting in concert on cell division but often pulling in opposite directions when it comes to branching (Romanov, 2022). The relevance of exogenous auxin is not limited to spray applications, either — chickpea rhizosphere bacteria that produce IAA have themselves been shown to promote growth (Lata et al., 2024), and auxin’s role in breaking seed dormancy and driving germination vigor, including through IAA-producing rhizobacteria, is an increasingly active area of legume agronomy (Ali and Moon, 2025). Recent screening work across dryland chickpea genotypes continues to underline how strongly genotype and management interact under these conditions (Maleki et al., 2024), and foliar trials in other semi-arid settings have reported real gains in nodulation, branching, and seed yield from inputs applied around flowering and pod set (Nair et al., 2024; Sridhara et al., 2024).

Cytokinin and auxin interactions extend to reproductive development as well: rather than acting independently, the two hormones appear to coordinate the initiation of floral and reproductive organs jointly, a mechanism with direct bearing on the flower-number and pod-set responses examined later in this paper (Baral et al., 2025). Cytokinin in particular has a documented role inside the developing seed itself, shaping the biosynthesis and translocation pathways that influence final seed size in cereals and legumes alike (Jameson, 2023) — relevant background for the grain-weight results discussed in Section 3.7.

This study set out to compare the separate and combined effects of foliar IAA and kinetin (0, 10, and 20 mg/L) on FLIP 84-92 and ILC 32-79 grown under two very different conditions — in the field at El Khroub and in pots at Oum El Bouaghi — during the 2024/2025 season. Three things motivated the design: describing how each growth and yield trait responded numerically to the nine hormone treatments at each site, comparing the two varieties against each other, and looking at how field and pot responses diverged. The data available to us were treatment means rather than individual replicate observations, so we stop short of claiming any statistical interaction or synergy.

## 2. Materials and Methods

### 2.1 Study Sites, Climate, and Soil

Field trials ran at the ITGC experimental station of El Khroub, roughly 14 km south of Constantine, while a parallel pot trial using the same two varieties was set up at the Institute of Natural Sciences in Oum El Bouaghi. Both trials covered the same window, from sowing in January 2025 through harvest in June 2025. We logged monthly minimum and maximum temperatures and total rainfall at each site from September 2024 through June 2025 (Table 1). Both sites saw their wettest stretch between winter and spring, though El Khroub received noticeably more rain overall (440.9 mm) than Oum El Bouaghi (326.7 mm) across the trial period.

**Table 1.**
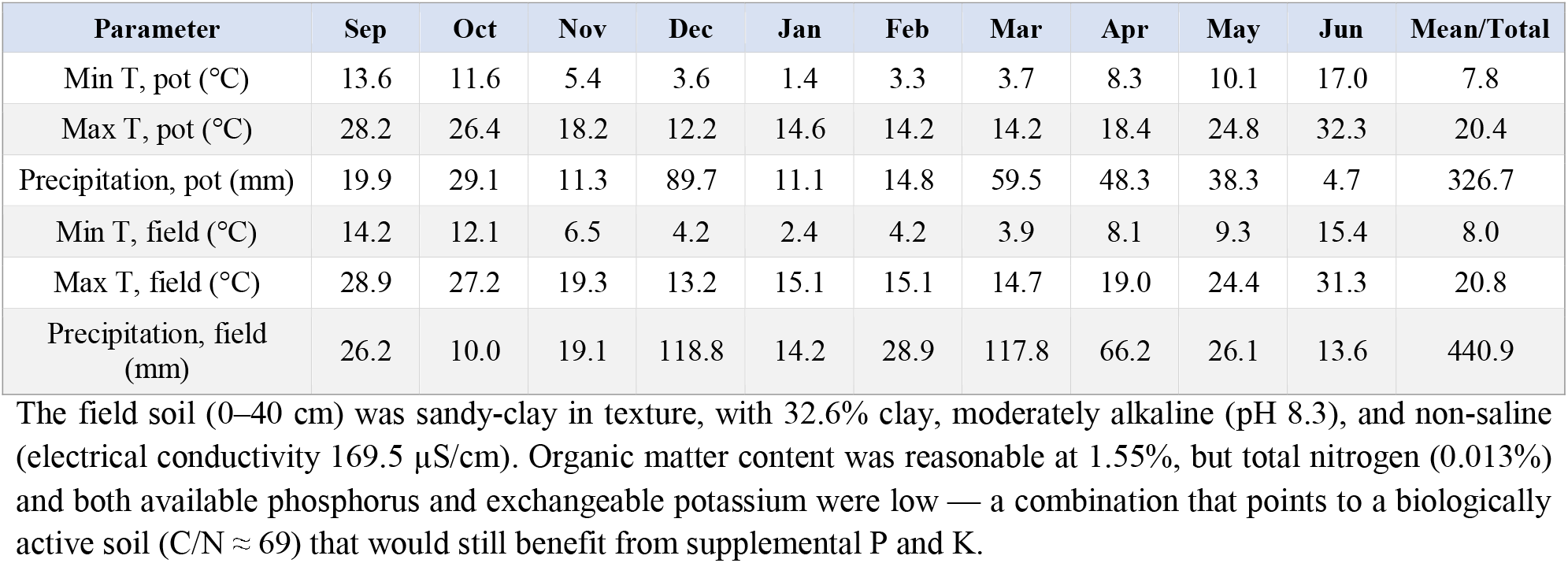
Monthly minimum/maximum temperature and precipitation at both sites, September 2024–June 2025.

| Parameter | Sep | Oct | Nov | Dec | Jan | Feb | Mar | Apr | May | Jun | Mean/Total |
| --- | --- | --- | --- | --- | --- | --- | --- | --- | --- | --- | --- |
| Min T, pot (°C) | 13.6 | 11.6 | 5.4 | 3.6 | 1.4 | 3.3 | 3.7 | 8.3 | 10.1 | 17.0 | 7.8 |
| Max T, pot (°C) | 28.2 | 26.4 | 18.2 | 12.2 | 14.6 | 14.2 | 14.2 | 18.4 | 24.8 | 32.3 | 20.4 |
| Precipitation, pot (mm) | 19.9 | 29.1 | 11.3 | 89.7 | 11.1 | 14.8 | 59.5 | 48.3 | 38.3 | 4.7 | 326.7 |
| Min T, field (°C) | 14.2 | 12.1 | 6.5 | 4.2 | 2.4 | 4.2 | 3.9 | 8.1 | 9.3 | 15.4 | 8.0 |
| Max T, field (°C) | 28.9 | 27.2 | 19.3 | 13.2 | 15.1 | 15.1 | 14.7 | 19.0 | 24.4 | 31.3 | 20.8 |
| Precipitation, field (mm) | 26.2 | 10.0 | 19.1 | 118.8 | 14.2 | 28.9 | 117.8 | 66.2 | 26.1 | 13.6 | 440.9 |

**Table 2.** Stem length (LT1, LT2), all nine hormone treatments, both sites.

| Treatment | Pot LT1 (cm) | Pot LT2 (cm) | Field LT1 (cm) | Field LT2 (cm) |
| --- | --- | --- | --- | --- |
| Control | 28.99 | 35.94 | 62.66 | 70.66 |
| A10 | 34.06 | 43.20 | 68.95 | 73.52 |
| A20 | 37.23 | 46.54 | 71.70 | 74.49 |
| K10 | 34.89 | 36.85 | 69.70 | 71.79 |
| K20 | 34.46 | 44.13 | 71.70 | 72.91 |
| A10+K10 | 30.06 | 45.97 | 72.29 | 75.70 |
| A10+K20 | 31.28 | 36.44 | 68.50 | 73.71 |
| A20+K10 | 33.36 | 36.22 | 64.37 | 74.05 |
| A20+K20 | 30.61 | 36.06 | 69.20 | 72.66 |

**Table 3.** Branching — primary and secondary branch number (NRP, NRS1, NRS2), all nine hormone treatments, both sites.

| Treatment | Field NRP | Pot NRS1 | Field NRS1 | Pot NRS2 | Field NRS2 |
| --- | --- | --- | --- | --- | --- |
| Control | 3.25 | 3.50 | 12.30 | 5.66 | 15.50 |
| A10 | 3.49 | 3.69 | 14.52 | 5.94 | 17.04 |
| A20 | 3.58 | 4.12 | 15.45 | 7.10 | 17.12 |
| K10 | 3.62 | 4.05 | 15.46 | 9.05 | 18.25 |
| K20 | 3.91 | 4.28 | 17.16 | 9.28 | 20.00 |
| A10+K10 | 3.58 | 4.27 | 13.93 | 7.59 | 18.12 |
| A10+K20 | 3.70 | 4.93 | 16.14 | 7.62 | 19.54 |
| A20+K10 | 3.62 | 4.73 | 16.12 | 8.04 | 18.25 |
| A20+K20 | 3.25 | 5.31 | 15.09 | 8.46 | 16.54 |

**Table 4.** Leaf number (NF1, NF2), all nine hormone treatments, both sites.

| Treatment | Pot NF1 | Field NF1 | Pot NF2 | Field NF2 |
| --- | --- | --- | --- | --- |
| Control | 29.50 | 67.50 | 88.75 | 95.00 |
| A10 | 64.42 | 82.08 | 89.25 | 120.00 |
| A20 | 57.66 | 85.83 | 89.00 | 134.00 |
| K10 | 62.94 | 90.00 | 91.16 | 113.50 |
| K20 | 69.83 | 115.41 | 94.12 | 138.08 |
| A10+K10 | 60.08 | 90.00 | 89.08 | 120.00 |
| A10+K20 | 57.16 | 105.00 | 94.80 | 132.83 |
| A20+K10 | 63.74 | 104.99 | 93.86 | 146.50 |
| A20+K20 | 63.41 | 77.91 | 95.83 | 137.50 |

**Table 5.**
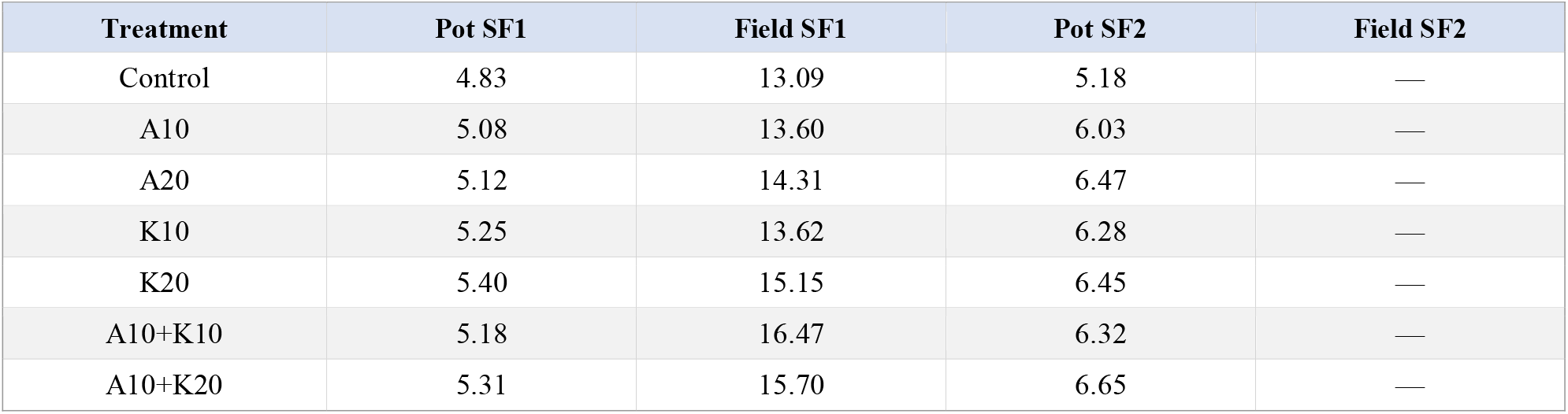
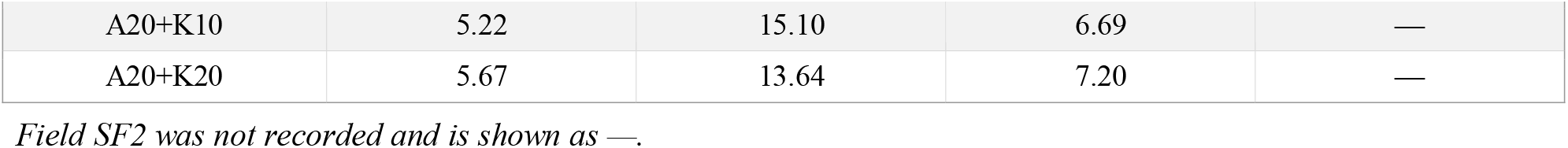
Leaf area (SF1, SF2, cm^2^), all nine hormone treatments, both sites.

**Table 6.**
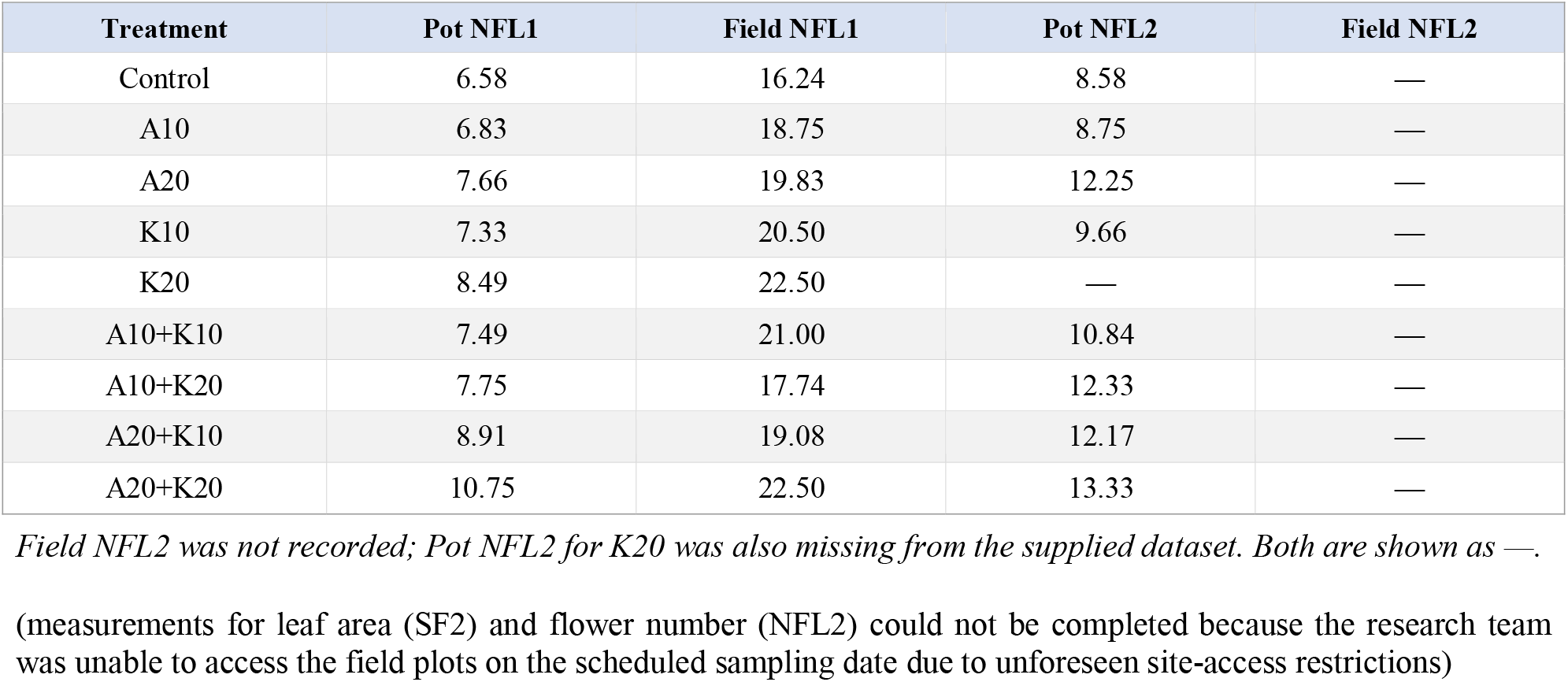
Flower number (NFL1, NFL2), all nine hormone treatments, both sites.

**Table 7.** Pod traits and dry matter weight, all nine hormone treatments, both sites.

| Treatment | Pot pod length (cm) | Field pod no. | Pot PMS (g) | Field PMS (g) |
| --- | --- | --- | --- | --- |
| Control | 2.12 | 9.50 | 4.16 | 61.62 |
| A10 | 2.30 | 15.16 | 5.17 | 62.70 |
| A20 | 2.21 | 16.93 | 5.42 | 68.19 |
| K10 | 1.99 | 18.30 | 4.27 | 56.92 |
| K20 | 2.20 | 21.28 | 5.29 | 71.45 |
| A10+K10 | 2.39 | 14.05 | 6.12 | 77.02 |
| A10+K20 | 2.38 | 14.44 | 4.35 | 84.97 |
| A20+K10 | 2.26 | 15.97 | 4.31 | 83.89 |
| A20+K20 | 2.42 | 14.60 | 5.38 | 95.21 |

**Table 8.**
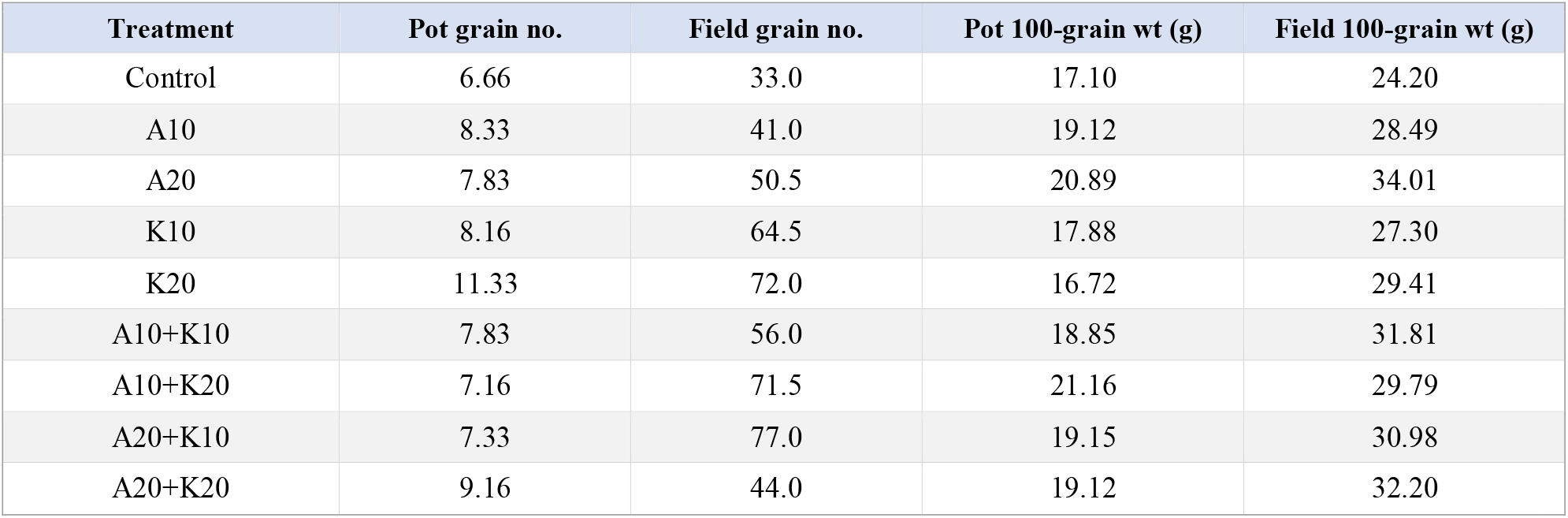
Grain number and 100-grain weight, all nine hormone treatments, both sites.

| Treatment | Pot grain no. | Field grain no. | Pot 100-grain wt (g) | Field 100-grain wt (g) |
| --- | --- | --- | --- | --- |
| Control | 6.66 | 33.0 | 17.10 | 24.20 |
| A10 | 8.33 | 41.0 | 19.12 | 28.49 |
| A20 | 7.83 | 50.5 | 20.89 | 34.01 |
| K10 | 8.16 | 64.5 | 17.88 | 27.30 |
| K20 | 11.33 | 72.0 | 16.72 | 29.41 |
| A10+K10 | 7.83 | 56.0 | 18.85 | 31.81 |
| A10+K20 | 7.16 | 71.5 | 21.16 | 29.79 |
| A20+K10 | 7.33 | 77.0 | 19.15 | 30.98 |
| A20+K20 | 9.16 | 44.0 | 19.12 | 32.20 |

### 2.2 Plant Material and Experimental Design

We worked with two chickpea varieties: FLIP 84-92, of Syrian origin, and ILC 32-79. In germination tests FLIP 84-92 came out slightly ahead, at 86% versus 83% for ILC 32-79. Seeds at El Khroub were sown mechanically on 21 January 2025 — 5 m rows, four rows per plot, 30 cm between plants, 1 m plot width, nine plots per variety — after tillage to 30–45 cm depth and an application of triple superphosphate (46%). The pot trial at Oum El Bouaghi used the same two varieties, germinated first in Petri dishes and then transplanted.

Nine hormone treatments made up the experimental layout: an untreated control, IAA alone at two rates (A10, A20), kinetin alone at two rates (K10, K20), and the four possible IAA × kinetin combinations (A10+K10, A10+K20, A20+K10, A20+K20). Each treatment was applied with three replicates per variety per site (9 × 3 × 2 = 54 treatment–replicate–variety units at each site). A10+K20, to take one example, means the final spray solution carried 10 mg/L IAA and 20 mg/L kinetin. We did not have enough detail on the original randomization to confirm a proper hierarchical split-split-plot layout, so rather than force a formal interaction test onto an unverifiable design, we treat the nine hormone combinations here as a single treatment factor and report descriptive comparisons only.

### 2.3 Hormone Application and Measured Traits

IAA and kinetin stock solutions were sprayed onto the foliage at 10 or 20 mg/L, alone or combined, in two applications 15 days apart. Vegetative traits were recorded 15 days after each spray (the suffixes 1 and 2 mark the first and second spray, respectively): stem length (LT1, LT2), primary branch number (NRP), secondary branch number (NRS1, NRS2), leaf number (NF1, NF2), leaf area (SF1, SF2), flower number (NFL1, NFL2), pod length in the pot trial, pod number in the field trial, dry matter weight (PMS), grain number (NGr), and 100-grain weight (P100Gr). Two field measurements — SF2 and NFL2 — were missing from the dataset we received and are left out of the analysis rather than estimated.

### 2.4 Statistical Reporting

A note on what kind of analysis this is: the manuscript we worked from reports treatment means and states that three replicates were used per treatment, but the underlying replicate-level values and any measure of spread were not part of the dataset. We have therefore not calculated P-values, run multiple-comparison tests, or tested for an IAA × kinetin interaction anywhere in this revision — what follows describes numerical differences between treatment means, not statistically established effects. Anyone revisiting this work should track down the original replicate-level records and analyze them properly before treating any of these differences as significant, let alone synergistic.

## 3 Results and Discussion

### 3.1 Germination and Growing Conditions

FLIP 84-92 germinated somewhat better than ILC 32-79 (86% vs. 83%), a gap of three points consistent with slightly stronger inherent vigour in the FLIP line. Conditions at both sites were broadly favorable for chickpea over the season, with rainfall peaking between winter and spring (Table 1). The El Khroub soil profile — decent organic matter but short on phosphorus and potassium — suggests early vegetative growth there may have been somewhat nutrient-limited despite otherwise workable conditions.

### 3.2 Stem Length (LT1, LT2)

Stem length responded differently at the two sites. At Oum El Bouaghi, IAA alone at the higher rate (A20) gave the tallest plants after both sprays (37.23 and 46.54 cm), with A10+K10 close behind after the second spray (45.97 cm). At El Khroub, by contrast, A10+K10 was the tallest treatment at both time points (72.29 and 75.70 cm). These are treatment averages, and without knowing how much individual plants varied within each treatment, we cannot say whether these gaps are meaningful.

### 3.3 Branching (NRP, NRS1, NRS2)

Primary branching was only recorded at El Khroub, where K20 produced the most branches (3.91), just ahead of A10+K20 (3.70); the control and A20+K20 tied for the fewest (3.25). Secondary branching followed a similar kinetin-favoring pattern at El Khroub — K20 topped both spray dates (17.16 and 20.00) — but at Oum El Bouaghi the picture split: A20+K20 led after the first spray (5.31) and K20 after the second (9.28). Kinetin-containing treatments turn up at or near the top of the branching tables fairly consistently, though again, we are looking at averages rather than tested differences.

### 3.4 Leaf Number and Leaf Area (NF1, NF2, SF1, SF2)

Leaf number tracked kinetin closely. K20 produced the most leaves at both El Khroub (115.41) and Oum El Bouaghi (69.83) after the first spray, though by the second spray at El Khroub, A20+K10 had overtaken it (146.50). Leaf area told a more site-specific story: A20+K20 was largest at Oum El Bouaghi at both sampling points (5.67 and 7.20 cm^2^), while A10+K10 led at El Khroub after the first spray (16.47 cm^2^). The second field leaf-area reading (SF2) was not recorded, so we cannot say anything about how that trait developed later in the season at El Khroub.

### 3.5 Flower Number (NFL1, NFL2)

In pots, A20+K20 produced the most flowers by the second spray (13.33). At El Khroub, K20 and A20+K20 tied for the highest first-spray count (22.50 each). This pattern fits what is known about auxin and cytokinin’s roles in growth and reproduction generally, but it does not on its own demonstrate that the two hormones interact statistically — that would need the replicate data we do not have. Field NFL2 was not recorded and is left out here as well.

### 3.6 Pod Traits and Dry Matter Weight

Pod number, recorded only at El Khroub, was more than double in the K20 treatment (21.28) compared with the untreated control (9.50). Dry matter weight peaked with A20+K20 in the field (95.21 g) and with A10+K10 in pots (6.12 g) — figures that are not directly comparable given how different the sampling basis and growing conditions were between the two trials, and that we cannot fully reconstruct from what is documented. What we can say is that the treatment associated with the highest biomass was not the same in both environments.

### 3.7 Grain Number and 100-Grain Weight

At Oum El Bouaghi, K20 gave the most grains per plant (11.33), while A10+K20 produced the heaviest 100-grain weight (21.16 g), with A20 close behind (20.89 g). At El Khroub the ranking shifted again: A20+K10 gave the most grains (77) and A20 the heaviest grains (34.01 g). Whichever treatment came out on top depended heavily on both the trait in question and the site. This kind of trait-by-site variability is consistent with auxin’s broader role in partitioning carbohydrate between vegetative and reproductive sinks, a mechanism demonstrated directly in rice (Zhao et al., 2022) and plausibly at work here too, though confirming it would need a proper factorial analysis with real replicate variance rather than the treatment means available to us.

**Figure 1.**
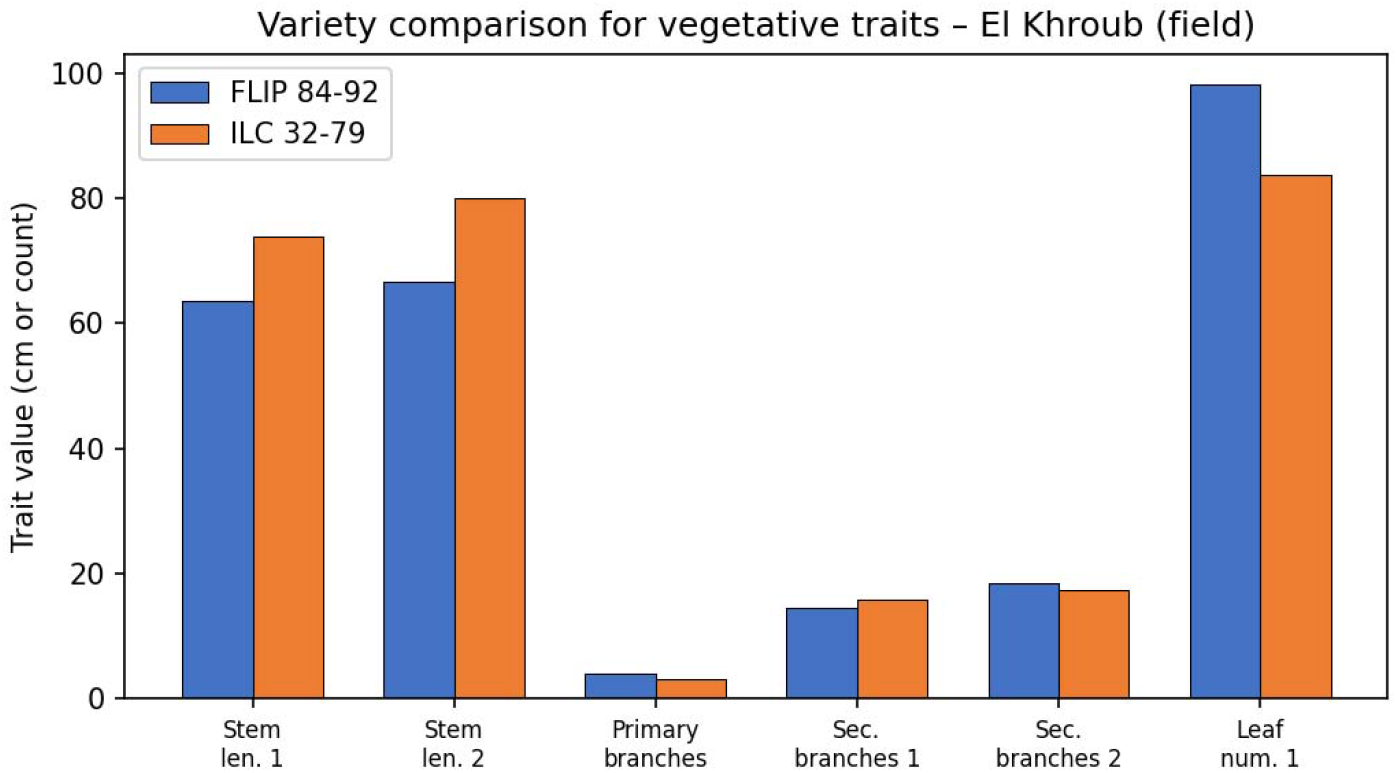
Variety comparison (FLIP 84-92 vs. ILC 32-79) for six vegetative traits, El Khroub (field trial).

### 3.8 Variety Comparison

Set side by side, the two varieties diverged mainly in stem length and leaf number rather than in branching. ILC 32-79 grew visibly taller stems than FLIP 84-92 at both sampling dates, while FLIP 84-92 carried more leaves after the first spray. Branch counts, both primary and secondary, were broadly similar between the two. Combined with the germination and grain-weight differences noted earlier, FLIP 84-92 comes across as the more productive of the two varieties under these conditions, while ILC 32-79’s height advantage may simply reflect a different growth habit rather than a yield benefit.

### 3.9 Field vs. Pot Comparison

**Figure 2.**
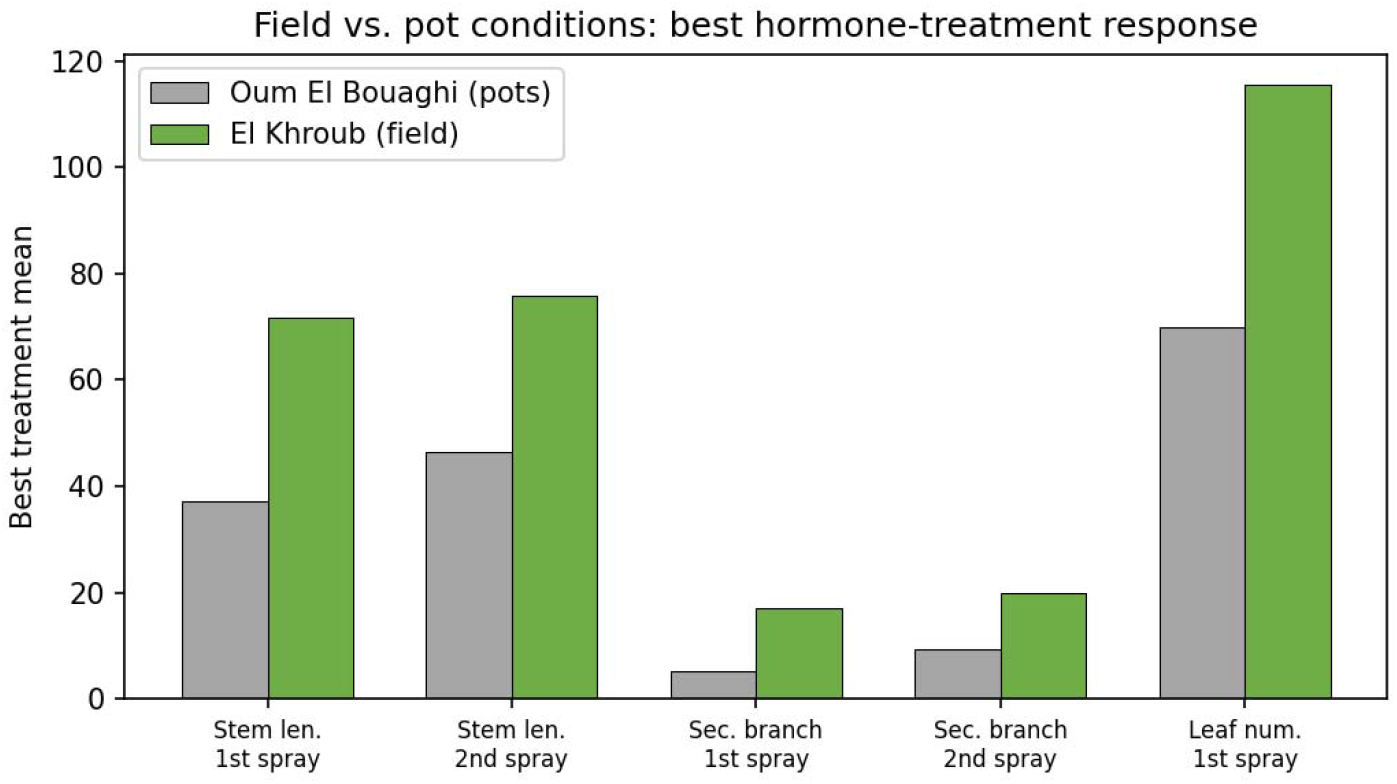
Field vs. pot comparison: best hormone-treatment response for five vegetative traits common to both trials.

No single hormone regime came out on top across every trait or both environments. Combined treatments accounted for some of the highest means — leaf area, flower number, dry matter weight, 100-grain weight — but kinetin alone was just as often the top performer, particularly for branching, leaf number, pod number, and grain number. K20 alone gave the most field pods and the most first-spray field leaves, for instance, while A20+K20 gave the largest pot leaf area and the highest field dry matter weight. None of this rules out an auxin–cytokinin interaction, but it does not confirm one either.

Field means ran consistently higher than pot means for most growth and yield traits, biomass especially. It is tempting to blame this on restricted root growth or weaker nodulation in pots, but we cannot actually attribute the gap to either mechanism here, since root growth, nodule counts, nitrogen fixation, and irrigation regime were not measured or documented in enough detail. Differences in rooting volume, water availability, and nutrient supply are all plausible contributors, but they remain hypotheses until someone measures them directly; work in other legumes has shown that nodulation itself is sensitive to fairly simple management choices — crushing nodules onto seed before sowing, for instance, measurably changes nodulation outcomes (Pudasaini et al., 2023) — which underlines how worthwhile it would be to measure this directly in a follow-up trial rather than assume it.

## 4. Conclusion

Foliar IAA and kinetin clearly moved the numbers for both chickpea varieties under the semi-arid conditions tested here, but which treatment came out on top depended on the trait and the environment in question. A20+K20 gave the largest pot leaf area and field dry matter weight; K20 alone led on field pod number and several leaf and branching measures; A10+K20 gave the heaviest pot grains. FLIP 84-92 germinated better and produced heavier grains than ILC 32-79, which in turn grew taller stems. Field responses ran well ahead of pot responses in absolute terms, though we cannot pin that gap on a specific root or nodulation mechanism with the data at hand.

## Data availability note

The tables here present treatment and variety mean only. Two field measurements, SF2 and NFL2, were missing from the supplied dataset and are flagged as such rather than estimated. The loss of these data points was due to logistical constraints during field sampling, not to analytical exclusion.

## Conflict of Interest

All authors declare no conflict of interest

## Ethics Approval

Not applicable to this paper

## Notes

### Competing Interest Statement

The authors have declared no competing interest.

